# Nanosized extracellular vesicles released by *Neurospora crassa* hyphae

**DOI:** 10.1101/2022.11.01.514727

**Authors:** Elizabeth Medina-Castellanos, Daniel A. Salgado-Bautista, Juan Manuel Martínez-Andrade, Ruben Dario Cadena-Nava, Meritxell Riquelme

## Abstract

Extracellular vesicles (EVs) are nanosized structures containing proteins, lipids, and nucleic acids, released by living cells to the surrounding medium. EVs participate in diverse processes, such as intercellular communication, virulence, and disease. In pathogenic fungi, EVs carry enzymes that allow them to invade the host or undergo environmental adaptation successfully. In *Neurospora crassa*, a non-pathogenic filamentous fungus widely used as a model organism, the vesicle-dependent secretory mechanisms that lead to polarized growth are well studied. In contrast, biosynthesis of EVs in this fungus has been practically unexplored. In the present work, we analyzed *N. crassa* culture’s supernatant for the presence of EVs by dynamic light scattering (DLS), transmission electron microscopy (TEM) and proteomic analysis. We identified spherical membranous structures, with a predominant subpopulation averaging a hydrodynamic diameter (*d*_*h*_) of 68 nm and a particle diameter (*d*_*p*_) of 38 nm. EV samples stained with osmium tetroxide vapors were better resolved than those stained with uranyl acetate. Mass spectrometry analysis identified 252 proteins, including enzymes involved in carbohydrate metabolic processes, oxidative stress response, cell wall organization/remodeling, and circadian clock-regulated proteins. Some of these proteins have been previously reported in exosomes from human cells or in EVs of other fungi. In view of the results, it is suggested a putative role for EVs in cell wall biosynthesis and vegetative development in *N. crassa*.

## 1. Introduction

Extracellular vesicles (EVs) are nanometric structures enclosed by a lipid bilayer produced and released by cells from the three domains of life. EVs production was initially regarded as a process to eliminate waste compounds from the cells (Johnstone et al., 1987). However, it was later revealed that EVs are associated with various biological processes, including cell-to-cell communication, host-pathogen interactions, and deadly human diseases, including cancer (Kalluri and LeBleu, 2020; Margolis and Sadovsky, 2019; Samuel et al., 2015). EVs are excellent indicators of physiological changes in a free-living cell or an entire tissue/organ as part of their interactions with the environment. Therefore, EVs’ proteins and nucleic acids are valuable biomarkers for disease diagnosis and health indicators (Hill, 2019; Lin et al., 2015). EVs comprise a large family of vesicles, which includes microvesicles or ectosomes (∼ 50-1000 nm), apoptotic bodies (500-1000 nm), and exosomes (∼ 40-160 nm) (Cocucci and Meldolesi, 2015; Kalluri and LeBleu, 2020). EVs’ size, content, type, and concentration may differ for each organism and for a specific physiological condition (Zhang et al., 2019). Exosome biogenesis begins with inward budding of the membrane of the late endosomes resulting in intraluminal vesicles (ILVs) within multivesicular bodies (MVBs). The membrane of the MVBs can fuse with the plasma membrane (PM), releasing the enclosed ILVs (which become the exosomes) to the extracellular space (Bebelman et al., 2018). In plant and animal cells, the ILVs formation is mediated by endosomal sorting complexes required for transport (ESCRT), sphingomyelinase 2 (nSMase2), and Syndecan-syntenin (Baietti et al., 2012; Bebelman et al., 2018). Kinesin and the Ras-related protein GTPase Rab7 mediate the transport of MVBs, whereas Rab27 and branched actin control the docking of MVBs with the PM (Kalluri and LeBleu, 2020; Raposo and Stoorvogel, 2013). In exosomes secreted by human cells, some commonly found marker proteins include tetraspanin, annexin, ESCRT proteins and their accessory proteins such as TSG101 (tumor susceptibility gene 101), ALIX (apoptosis-linked gene 2-interacting protein X), HSC70 (heat shock cognate 70), and HSP90β (heat shock protein 90) (Doyle and Wang, 2019).

In fungi, the biogenesis and role of EVs have been largely underexplored. The first studies on fungal EVs were carried out in the basidiomycete human pathogenic yeast *Cryptococcus neoformans*. This fungus releases EVs that transport its major capsular polysaccharide glucuronoxylomannan to the extracellular space (Rodrigues et al., 2007; Yoneda and Doering, 2006). The biological roles of *C. neoformans* EVs in its interaction with host cells have been extensively studied (de Oliveira et al., 2020). A growing number of studies have identified EVs in other fungal species, including the human pathogens *Aspergillus flavus* (Brauer et al., 2020), *A. fumigatus* (Souza et al., 2019), *Paracoccidioides brasiliensis* (Peres da Silva et al., 2019; Vallejo et al., 2011), and *Candida albicans* (Albuquerque et al., 2008; Peres da Silva et al., 2015; Rodrigues et al., 2008b), as well as the non-pathogenic yeast *Saccharomyces cerevisiae* (Oliveira et al., 2010). Fungal EVs contain RNA molecules (Albuquerque et al., 2008; Peres da Silva et al., 2019; Peres da Silva et al., 2015; Rodrigues et al., 2008b). In *Histoplasma capsulatum* EVs contain components of the microRNA machinery, suggesting that human fungal pathogens can interfere with the host’s gene expression (Alves et al., 2019).

Yeast EVs have been classified as exosomes, given their similarities in cargo and morphology with mammalian exosomes (Casadevall et al., 2009; Panepinto et al., 2009; Rodrigues et al., 2008a). However, components of the non-conventional secretory pathway for exosomal biogenesis in mammals like Vps23 and ALIX are not present in fungal EVs (Bleackley et al., 2019). In addition, there is evidence that fungal EVs biogenesis involves components of the conventional post-Golgi secretory pathway, such as Sec1-1, Sec4, and Bos1-1 (Oliveira et al., 2010; Yoneda and Doering, 2006), which suggests that the synthesis of EVs is dependent on conventional (ER/Golgi-dependent), as well as unconventional pathways (ER/Golgi-independent) (Miura and Ueda, 2018; Oliveira et al., 2010). Given this, fungal EVs biogenesis could involve similar mechanisms to those participating in the biogenesis of secretory vesicles during hyphal growth but following distinct routes and transport mechanisms.

*Neurospora crassa*, an important model system for secretion and polarized growth, presents exocytic events at various sites within the hyphal apex during vegetative growth. Hyphal elongation in *N. crassa* involves the accumulation of secretory vesicles at the Spitzenkörper (SPK), an apical body that determines hyphal growth, direction, and morphology (Riquelme, 2013). A population of these vesicles is thought to associate with the exocyst complex before fusing with the PM, with the subsequent incorporation of transmembrane proteins (Riquelme et al., 2014). Remarkably, chitin and *β*-1,3-glucan synthase activities have been identified in the chitosomes (microvesicles) and macrovesicles, respectively (Sanchez-Leon et al., 2011; Verdin et al., 2009). Besides cell wall synthesis associated functions, secretory vesicles targeted to the apical zone presumably carry signals for cell fusion, including, for instance, a MAK-2 MAPK module that is recruited to the PM and alternates between sending and receiving signals during cell fusion events, a highly coordinated process in germlings (Fleissner et al., 2009) and vegetative hyphal fusion in *N. crassa*. (Serrano et al., 2018). *N. crassa* presents MVBs diverse in size and shape, which are found in subapical and distal hyphal regions (beyond 30 to 40 µm from the tip), as seen by transmission electron microscopy (TEM) (Rico-Ramírez et al., 2018), suggesting that it is also capable of producing exosomal EVs. However, no studies show the fusion of MVBs with the PM and subsequent release of EVs in *N. crassa* or other filamentous fungi.

In this study, we processed and analyzed *N. crassa* culture supernatants and characterized the isolated EVs by dynamic light scattering (DLS), TEM, and liquid chromatography with tandem mass spectrometry (LC-MS/MS). We identified a predominant subpopulation of EVs similar in morphology and size to that reported for exosomes. Furthermore, among all the proteins found as EV cargoes, there was a prevalence of proteins related to cell wall remodeling processes and vesicular trafficking.

## 2. Materials and methods

### 2.1. Fungal strains and growth conditions

*N. crassa* N1 strain (FGSC# 988) was grown in Vogel’s minimal medium (VMM) supplemented with 1.5% sucrose (Vogel, 1956). For conidia production, the mycelium was grown on flasks containing solid VMM at 30 °C for 72 h in the dark, and then exposed to the light at room temperature for additional 48 h. Finally, the conidia were collected in 1M sorbitol solution and stored at -20 °C.

### 2.2. Isolation of extracellular vesicles

Conidia of *N. crassa* N1 strain (1 × 10^8^) were inoculated into 200 ml of liquid VMM and incubated in an orbital shaker (150 rpm) at 30 °C for 48 h. EDTA (10 mM) was added to the culture at the end of the incubation time to avoid aggregates. The mycelium was filtered out by using Whatman® filter paper #1 (11 μm pore size). To obtain EVs from the resulting filtrate a differential ultracentrifugation method (Rodrigues *et al*., 2007) with slight modifications was carried out. The supernatant was centrifuged sequentially at 3 000 rpm for 30 min to remove cell debris and apoptotic bodies and at 10 000 rpm for 1 h to eliminate large vesicles. The final supernatant was ultracentrifuged (Optima MAX-XP Beckman Coulter) at 100 000 x g (MLA-50 fixed angle rotor) for 2 h, and the resulting pellet was resuspended with 200 μL of 20 mM Tris-HCl (pH 7.2) for DLS and TEM analyses, or with 200 μL of Exosome Resuspension Buffer (1-2.5% Triton X-100) from the Total Exosome RNA and Protein Isolation Kit (Invitrogen, Thermo-Fisher Scientific) to lyse the EVs for proteomic analysis.

The dry weight of the biomass was estimated gravimetrically, after drying the mycelium for 24 h at 60 °C.

### 2.3. Preparation of catanionic vesicles

SDS-CTAB vesicles were prepared as previously described (Tah et al., 2011; Tah et al., 2014) with slight modifications. A stock solution of the anionic surfactant Sodium Dodecyl Sulfate (SDS) was prepared at a concentration of 1 mM in 1:1 chloroform and methanol solvent, and the cationic surfactant Cetyl Trimethyl Ammonium Bromide (CTAB) was prepared at a concentration of 1 mM in chloroform. Surfactant stock solutions were mixed at a 35:65 volume ratio of SDS:CTAB to obtain the catanionic vesicles. Then, prepared surfactant solution in organic solvent was heated in a water bath at 50 °C to evaporate the solvent in an atmosphere of nitrogen. Subsequently, phosphate buffered saline (PBS) was added to form vesicles. Immediately, the solution was sonicated for 10 min using a water bath sonicator (Fisher Scientific) to form vesicles. Next, the solution was extruded 10 times through a 0.22 µm Whatman filter attached to a syringe, and the vesicles were extruded 10 times again with a 0.1 µm filter (Whatman) to obtain small vesicles. The SDS-CTAB catanionic vesicles were stored at 4°C and used as controls in TEM analyses.

### 2.4. Dynamic light scattering analysis

The hydrodynamic diameter (*d*_*h*_) of EV preparations was measured by dynamic light scattering (DLS) using a Zetasizer Nano ZS equipment (ZEN3600, Malvern Instruments, Malvern, UK). One hundred microliters of the suspension of purified EVs in 20 mM Tris-HCl (pH 7.2) from *N. crassa* were analyzed using a disposable plastic micro cuvette (Malvern Panalytical Inc., ZEN0040). The measurements were done by triplicate (9 measurements with automatic runs (12-16 runs)) and analyzed using NanoSight software (version 3.2.16).

### 2.5. Transmission Electron Microscopy

EVs were stored at -20°C until processing. For each staining method, 10 *μ*L of each of the EVs samples were dropped in 200 mesh formvar-carbon copper grids (Ted Pella, Inc) and were let stand for 1 min. After that, the excess of liquid was removed with blotting paper. Immediately after, the samples were stained as follows: 1) for uranyl acetate staining, a fresh aqueous solution of 2% uranyl acetate (w/v) was prepared and centrifuged for 10 min at 11,000 rpm before being used. Ten µL of the uranyl acetate solution were dropped onto the grid, let stand for 60 s, and then blotted away carefully; 2) for osmium tetroxide vapors staining, crystals of osmium tetroxide were dissolved in an aqueous solution at 2% (w/v) before use. The grids were held with tweezers 2-5 cm above the lids of 1.5 mL Eppendorf tubes containing the osmium tetroxide solution for at least 10 min in a fume hood. Observations were conducted in a Hitachi H7500 transmission electron microscope (80 kV) equipped with a 16-megapixel Gatan CCD digital camera. The particle diameters (*d*_*p*_) were manually measured using ImageJ (NIH). The software was calibrated with the bar of the images obtained by TEM.

### 2.6. Protein separation by SDS-PAGE and tryptic digestion

The concentration of protein from EVs was quantified with a BCA Protein Assay Kit (23225, Thermo-Fisher Scientific). EV samples were aliquoted into 25 µg, and cleaned with the ReadyPrep 2-D cleanup kit (1632130, Bio-Rad). The pellet obtained was dissolved in 10 μL of 20 mM Tris-HCl (pH 7.2) and 10 μL of Laemmli sample buffer (Mini-PROTEAN® Tetra Cell-Bio-Rad). The solubilized proteins were heated (95 °C, 4 min) and centrifuged (10,000 rpm, 10 min) to eliminate suspended solids. Finally, the denaturalized proteins were separated in a 12% SDS-PAGE precast gel (Mini-PROTEAN TGX Precast Gels, Bio-Rad). Electrophoresis was run at 250 V (PowerPac Basic power supply, Bio-Rad) until the running front of the Laemmli buffer reached the end of the gel. Protein bands were visualized by staining with Coomassie Brilliant Blue G-250 (161–0406, Bio-Rad).

For Gel-LC analysis, each gel lane was cut into different regions (specific molecular weight regions with low abundance of proteins and regions with high abundance of proteins) to resolve possible issues such as dynamic range. The gel pieces were faded with 50 mM ammonium bicarbonate/50% acetonitrile followed by reduction with DTT and alkylation with iodoacetamide. The digestion was performed with trypsin at 37 °C overnight. The digested peptides were extracted from the gel twice using formic acid/acetonitrile/water (1:2:97) and water/acetonitrile (50:50), respectively. Finally, the tryptic digested peptides were concentrated in a centrifuge concentrator (SpeedVac, Thermo Savant) and kept to -80 °C until their analysis by LC-MS/MS.

### 2.7. Protein identification by liquid chromatography-tandem mass spectrometry analysis

Peptides separation was performed using an Ultimate 3000RS UHPLC in nanoflow configuration (Thermo Scientific, Bremen, Germany) coupled to QExactive Plus (Thermo Scientific, Bremen, Germany) quadrupole Orbitrap through a nanoelectrospray ion source, using Full MS followed by ddMS^2^ (DDA) during 110 min according to Kelstrup *et al*. (2012), Sun *et al*. (2013) and Liu *et al*. (2019). Mascot Distiller v2.6.2.0 in-house licensed (www.matrixscience.com) and Proteome Discoverer v2.5 (Thermo Scientific) were used to generate the peak list at the mascot generic format (mgf) to identify +1 or multiple charged precursor ions from the mass spectrometry data file. Parent mass (MS) and fragment mass (MS/MS) peak ranges were 250-1800 Da (resolution 70000) and 65-2000 Da (resolution 17500), respectively. Mascot server v2.8.1 (www.matrix-science.com, UK) in MS/MS ion search mode (local licenses) was applied to conduct peptide matches (peptide masses and sequence tags) and protein searches against *N. crassa* v20220513 from www.uniprot.org (24013 sequences, 11518915 residues). The following parameters were set for the search: carbamidomethyl (C) on cysteine was set as fixed; variable modifications included asparagine and glutamine deamidation and methionine oxidation. Only two missed cleavages were allowed; monoisotopic masses were counted; the precursor peptide mass tolerance was set at 20 ppm; fragment mass tolerance was 0.02 Da and the ion score or expected cut-off was set at 5. The MS/MS spectra were searched with MASCOT using a 95% confidence interval (C.I. %) threshold (p<0.05), with a minimum score of 22 used for peptide identification, indicating identity or extensive homology. Five lanes (replicas) were analyzed by LC-MS/MS.

### 2.8. Bioinformatic analysis of proteins

The characterization of identified proteins was done with different databases. Peptide signals were predicted with SignalP 6.0 and transmembrane regions were predicted with TMHMM 2.0. The rest of the information of Supplementary Table 1 was obtained directly with UniProt. In addition, the uncharacterized proteins were blasted in UniProt or NCBI blast service to obtain more information.

The classification of proteins according to the functional analysis was done with the Omicsbox software (version 2.1.14) (OmicsBox – bioinformatics made easy, BioBam Bioinformatics, March 3, 2019, from https://www.biobam.com/omicsbox). The sequences of all proteins were loaded in FASTA file format. To obtain the most complete annotation labels we used the Gene Ontology annotation workflow. The sequences were blasted using NCBI reference database, then the GO mapping was used to assign the most reliable GO terms to the input sequences considering the GO hierarchy. Graphs for the three categories (biological process, cellular component and molecular function) were combined as histograms that express direct counts (number of sequences). The relative abundance was determined by the emPAI value given by MASCOT. The total sum of emPAI values from the identified proteins represented one hundred percent.

## 3. Results

### 3.1. Characterization of EVs released by N. crassa

The *N. crassa* biomass recovered (dry weight) from the 200 mL culture used for EV analyses was 0.93 ± 0.05 g. DLS analysis of the cultures’ supernatants indicated presence of EVs of different sizes, with an average *d*_*h*_ of 68.048 ± 37.45 nm (range 24-141 nm), considering those particles with an abundance percentage greater than 1% (Fig. 1). This represents the 98.14% of the total particles measured by DLS. The large particles were later resolved by TEM as vesicle clusters of two or more units (Figure S1).

**Fig. 1.**
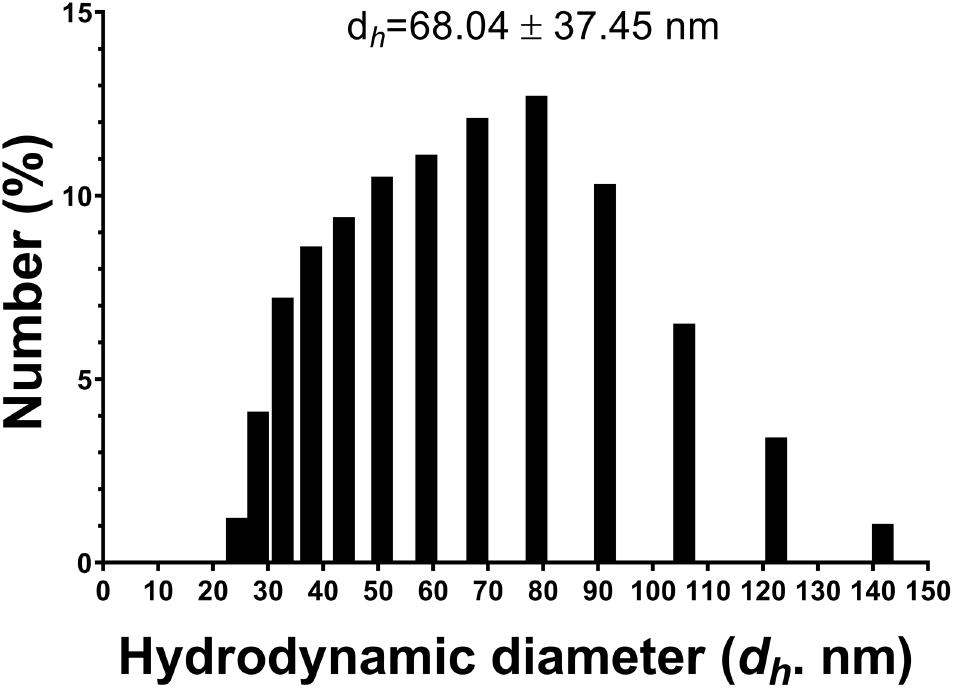
Dynamic light scattering of EVs from *N. crassa*. (n = 41). The average and standard deviation correspond to 98.14 % of the total measured particles.

### 3.2. Morphological characterization of N. crassa EVs

To determine the presence of EVs, the samples were imaged by TEM using two staining techniques, osmium tetroxide vapors and uranyl acetate. Spherical structures were observed by either method (Fig. 2A, B). Both SDS-CTAB vesicles and EVs had a lipidic membrane, which was disrupted upon treatment with Exosome Resuspension Buffer. The average *d*_*p*_ obtained was 33.65 ± 13.23 nm and 38.40 ± 15.50 nm for the osmium tetroxide vapors and the uranyl acetate method, respectively (Fig. 2C). There were no significant differences in the EVs *d*_*p*_ using either method of staining. However, in the samples treated with osmium tetroxide vapors there was not such a dark background, and EVs were resolved with a higher definition (Fig. 2A) than in the samples treated with uranyl acetate (Fig. 2B).

**Fig. 2.**
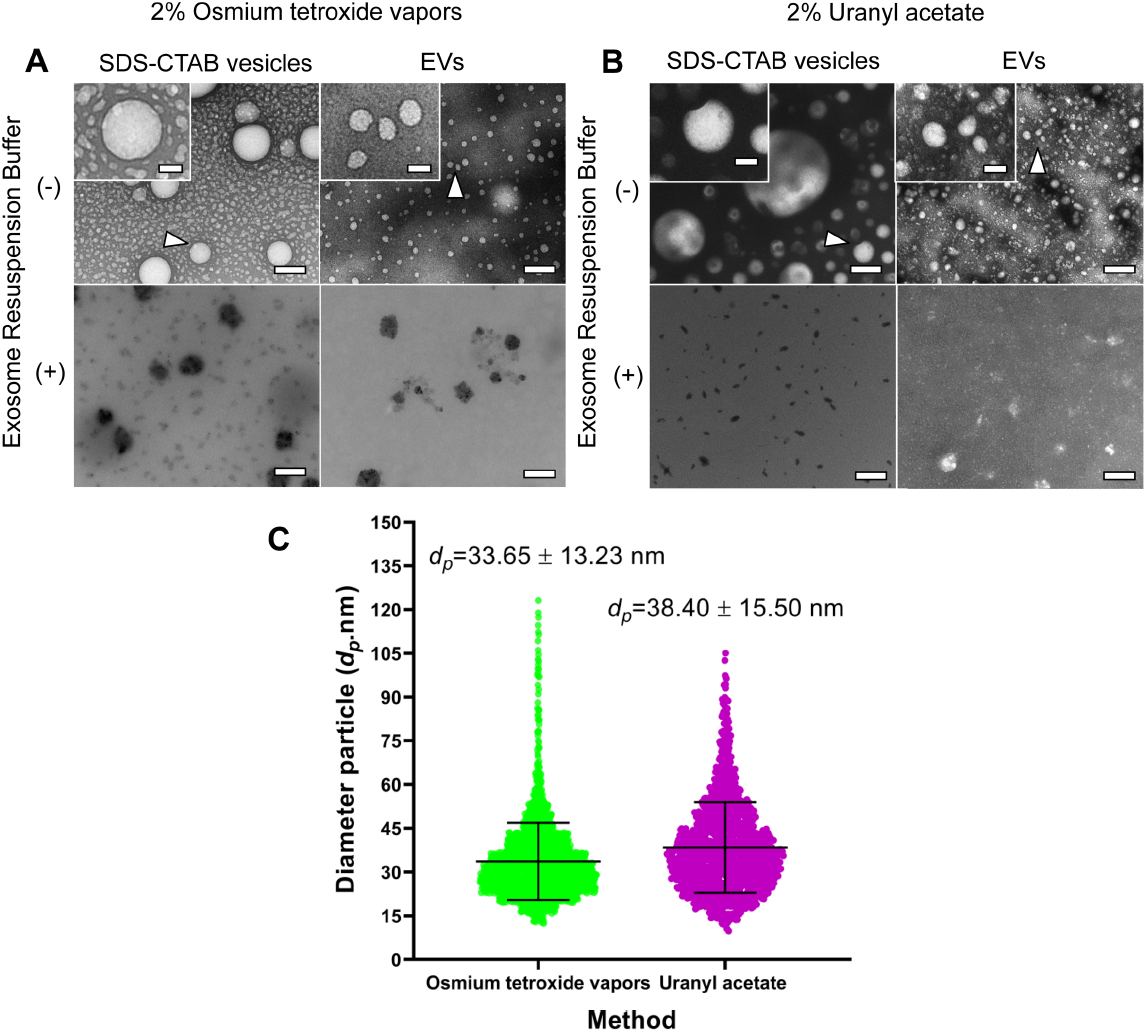
Characterization of EVs from *N. crassa* by transmission electron microscopy. A) Stained EVs with osmium tetroxide vapors and B) stained EVs with uranyl acetate. In both cases the membrane contrast was comparable to the SDS-CTAB vesicles used as control (+), although higher membrane contrast was notorious for the osmium tetroxide vapors. To confirm the membranous nature of the EVs, the samples were treated with Exosome resuspension buffer (ERB), which affected the presence of EV-like structures as well as of the SDS-CTAB vesicles. Bar in main box = 200 nm; bar in magnification box=50 nm. C) Measurements from images obtained with TEM using ImageJ. Osmium tetroxide vapors n=2068; uranyl acetate n= 1320.

Additionally, some EVs clusters were observed (Fig. S1). Their overall size was variable, depending on the number of EV units that they contained. Clusters of two or more units displayed a maximum diameter of 133 nm.

### 3.3. N. crassa EVs proteome contains diverse cellular functions

The total quantified protein obtained from pulled EV samples was 135.23 ± 23.55 µg. After separating 25 µg per lane by SDS-PAGE, proteins ranging from 10 to 150 kDa were selected (Fig. S3), and analyzed by LC-MS/MS (raw results can be found in Supplementary Table S1). In total, 252 proteins were identified as putative cargoes of *N. crassa* EVs (Supplementary Table S2). Of these, 48.8% of the proteins contain a signal peptide, and 11.5% of the proteins contain transmembrane domains. The emPAI value obtained by MASCOT indicates that 57 of the 252 proteins represent the 90% of the total emPAI value (Table 1) and 15 of these have emPAI values >1%, among which we can highlight the cell wall protein PhiA (NCU00399) and Clock controlled protein CCG-14 (NCU07787) with 26.26% and 19.20% emPAI values, respectively.

**Table 1.** Identified proteins from EVs of *N. crassa* that represent 90% of the total emPAI value.

| Entry UniProtKB | Gene names/Locus tag | L | # seq | % emPAI | Description |
| --- | --- | --- | --- | --- | --- |
| Q1K677 | *gel-3* NCU08909 | 542 | 18 | 0.61 | 1,3-beta-glucanosyltransferase (SP)* |
| F5HEQ0 | NCU08720 | 180 | 13 | 2.80 | AA1-like domain-containing protein (SP) |
| Q1K8A5 | NCU01063 | 208 | 7 | 0.46 | AA1-like domain-containing protein (SP) |
| A0A0B0EG60 | GE21DRAFT_2394 | 393 | 24 | 1.34 | Alpha/beta-hydrolase (SP) |
| A0A0B0DEM7 | GE21DRAFT_3704 | 342 | 9 | 0.33 | Alpha-L-arabinofuranosidase (SP) |
| Q7S4H9 | *acw-12* NCU08171 | 381 | 26 | 3.00 | Anchored cell wall protein 12 (SP) |
| Q7SAW7 | *acw-3* NCU05667 | 219 | 5 | 0.23 | Anchored cell wall protein 3 (SP) |
| Q1K686 | *acw-5* NCU07776 | 190 | 4 | 0.23 | Anchored cell wall protein 5 (SP) |
| V5IPM4 | NCU10852 | 628 | 23 | 0.32 | Beta-hexosaminidase (SP) |
| A0A0B0DXX7 | GE21DRAFT_5887 | 353 | 10 | 0.25 | Carbohydrate esterase family 16 protein (SP) |
| A0A0B0DWR4 | GE21DRAFT_5207 | 296 | 7 | 0.24 | Carbohydrate esterase family 5 protein (Fragment) (NSP) |
| Q1K6X1 | NCU06720 | 577 | 17 | 0.26 | Carboxypeptidase (SP) |
| A0A0B0EB54 | *cat-3* GE21DRAFT_4671 | 719 | 40 | 1.39 | Catalase (SP) |
| Q7RYR1 | NCU00399 | 198 | 12 | 26.26 | Cell wall protein PhiA (SP) |
| Q7S526 | NCU05882 | 422 | 14 | 0.94 | Cellulase (SP) |
| Q1K679 | *ccg-14* NCU07787 | 138 | 8 | 19.20 | Clock controlled gene CCG-14 (SP) |
| Q1K6S6 | *ccg-15* NCU08936 | 411 | 19 | 3.32 | Clock-controlled gene CCG-15 (SP) |
| Q6MVI0 | B23L4.170 | 1105 | 40 | 0.82 | Copper radical oxidase (SP) |
| A0A0B0E7T7 | *cyn-1* GE21DRAFT_7677 | 181 | 9 | 0.89 | Cyanate hydratase (NSP) |
| A0A0B0DFH2 | GE21DRAFT_3204 | 108 | 4 | 0.35 | Cytochrome c (NSP) |
| V5IQ13 | *scp* NCU00831 | 561 | 21 | 0.34 | Extracellular serine carboxypeptidase (SP) |
| A0A0B0DRB8 | GE21DRAFT_3513 | 407 | 15 | 0.28 | Galactose mutarotase-like protein (SP) |
| A0A0B0E9D9 | GE21DRAFT_3470 | 663 | 27 | 0.77 | Glucoamylase (NSP) |
| A0A0B0DS77 | GE21DRAFT_3270 | 594 | 18 | 0.44 | Glutathione hydrolase (SP) |
| A0A0B0E3C7 | GE21DRAFT_9367 | 364 | 16 | 0.63 | Glycosidase (SP) |
| Q6M9E3 | GE21DRAFT_6080 | 287 | 14 | 0.76 | Glycoside hydrolase family 16 protein (SP) |
| A0A0B0DDF8 | GE21DRAFT_10321 | 379 | 23 | 3.11 | Glycoside hydrolase family 28 protein (SP) |
| A0A0B0E8A2 | GE21DRAFT_6123 | 480 | 18 | 0.42 | Glycoside hydrolase family 30 protein (SP) |
| Q7S340 | *bgt-2* NCU09175 | 410 | 16 | 0.80 | GPI-anchored cell wall beta-1,3-endoglucanase EglC (SP) |
| A0A0B0EAA1 | GE21DRAFT_6010 | 172 | 7 | 1.00 | GPI-anchored domain-containing protein (SP) |
| A0A0B0DKP3 | GE21DRAFT_1437 | 257 | 7 | 0.23 | GPI-anchored domain-containing protein (SP) |
| A0A0B0E8G7 | GE21DRAFT_6027 | 211 | 11 | 1.50 | Hce2 domain-containing protein (SP) |

| Entry<br>UniProtKB | Gene names/Locus tag | L | # seq | % emPAI | Description |
| --- | --- | --- | --- | --- | --- |
| Q1K7I2 | *inv* NCU04265 | 695 | 29 | 1.31 | Invertase (SP) |
| A0A0B0EG67 | GE21DRAFT_2015 | 226 | 9 | 0.60 | Lysozyme (SP) |
| Q1K5A2 | *ncw-1* NCU05137 | 691 | 27 | 2.74 | Non-anchored cell wall protein 1 (SP) |
| Q7SET0 | *ncw-5* NCU00716 | 154 | 5 | 0.54 | Non-anchored cell wall protein 5 (SP) |
| Q7S4I2 | NCU02343 | 670 | 26 | 0.41 | Non-reducing end alpha-L-arabinofuranosidase (SP) |
| A0A0B0ED59 | GE21DRAFT_6132 | 177 | 9 | 0.54 | Phosphatidylglycerol/phosphatidylinositol transfer protein (SP) |
| Q1K5X6 | NCU07159 | 396 | 8 | 0.26 | Proteinase T (SP) |
| A0A0B0DXV1 | GE21DRAFT_6334 | 503 | 28 | 1.27 | Purple acid phosphatase (SP) |
| Q6MWR8 | B18P7.160 | 407 | 16 | 0.31 | Related to aldose 1-epimerase (SP) |
| Q6MVH1 | B20J13.080 | 215 | 6 | 0.52 | Related to extracellular matrix protein (SP) |
| A0A0B0E813 | GE21DRAFT_8917 | 540 | 20 | 0.43 | Rhamnogalacturonate lyase (SP*) |
| Q7S134 | NCU09992 | 547 | 22 | 0.57 | Serine peptidase (SP) |
| A0A0B0DNR4 | GE21DRAFT_3645 | 633 | 23 | 0.67 | Subtilisin-like protein (NSP) |
| Q9C1L1 |  | 107 | 7 | 1.32 | Thioredoxin (NSP) |
| A0A0B0DT50 | GE21DRAFT_289 | 324 | 14 | 0.43 | Transaldolase (NSP) |
| A0A0B0DNL9 | GE21DRAFT_4928 | 248 | 7 | 0.24 | Triosephosphate isomerase (NSP) |
| A0A0B0DRZ7 | GE21DRAFT_1379 | 120 | 5 | 0.29 | Tubulin-specific chaperone A (NSP) |
| P0C224 | *crp-79* NCU05275 | 128 | 8 | 0.76 | Ubiquitin-60S ribosomal protein L40 (NSP) |
| A0A0B0DSY8 | GE21DRAFT_9822 | 208 | 6 | 0.22 | Uncharacterized protein/Aldolase_II domain-containing protein (SP) |
| A0A0B0DEW9 | GE21DRAFT_1686 | 139 | 5 | 0.30 | Uncharacterized protein/Bac_luciferase domain-containing protein (SP) |
| A0A0B0DZG2 | GE21DRAFT_1911 | 312 | 8 | 0.33 | Uncharacterized protein/Candida_ALS_N domain-containing protein (NSP) |
| A0A0B0E6Q3 | GE21DRAFT_9470 | 120 | 6 | 1.03 | Uncharacterized protein/DNA repair protein RadA (SP) |
| Q7SBK9 | NCU08541 | 136 | 5 | 0.59 | Uncharacterized protein/NicO-domain-containing protein (NSP) |
| A0A0B0DL38 | GE21DRAFT_9087 | 178 | 5 | 0.24 | Uncharacterized protein/Putative phospholipase d1 protein (SP) |
| Q1K6F0 | NCU04675 | 221 | 8 | 0.53 | Uncharacterized protein/UPF0172-domain-containing (SP*) |

L=length; #seq= number of significant sequences; SP=signal peptide; NSP=No signal peptide; (\*) presence of transmembrane domains.

The proteins identified were classified by functional annotation with OmicsBox (Fig. 3, Supplementary Table S2) into the following groups: Biological processes (Fig. 3A), cellular component (Fig. 3B), and molecular function (Fig, 3C). The metabolic processes category was the most represented within the biological processes (Fig. 3A). This group contained a great number of proteins such as four isoforms of 1,3-*β*-glucanosyltransferases and diverse glycosidases from different families. The most abundant in this group were a glycoside hydrolase family 28 protein (A0A0B0DDF8), a non-anchored cell wall protein 1 (NCW-1; NCU05137), a catalase (CAT-3; GE21DRAFT_4671) and a purple acid phosphatase (GE21DRAFT_6334). Within the cellular component group, the most abundant proteins fall within the following categories: cellular anatomical entity (34 proteins), organelle (26 proteins), intracellular organelle (25 proteins), intracellular anatomical structure (25 proteins) and membrane-bounded organelle (25 proteins) (Fig. 3B). Some of the proteins identified in the cellular anatomical entity were two vacuolar protein sorting-associated proteins (GE21DRAFT_1332386 and NCU04137; VPS-5), and a coatomer subunit beta (GE21DRAFT_6111). Finally, within the molecular function group the categories that presented the highest number of proteins were the catalytic activity with 107 proteins, binding activity with 60 proteins, and hydrolase activity with 59 proteins (Fig. 3C). Hydrolases were some of the most abundant proteins. Glycoside hydrolases from distinct families and 1,3-*β*-glucanosyltransferases within the catalytic activity category were greatly represented in the biological processes and the molecular function groups. Interestingly, in the molecular function group we identified proteins involved in the endocytic trafficking, vesicular transport, and related with vacuolar membranes, such as phosphatidylinositol 3-kinase VPS-34 (GE21DRAFT_2066) and the vacuolar protein sorting-associated protein VPS-5 (NCU04137).

**Fig. 3.**
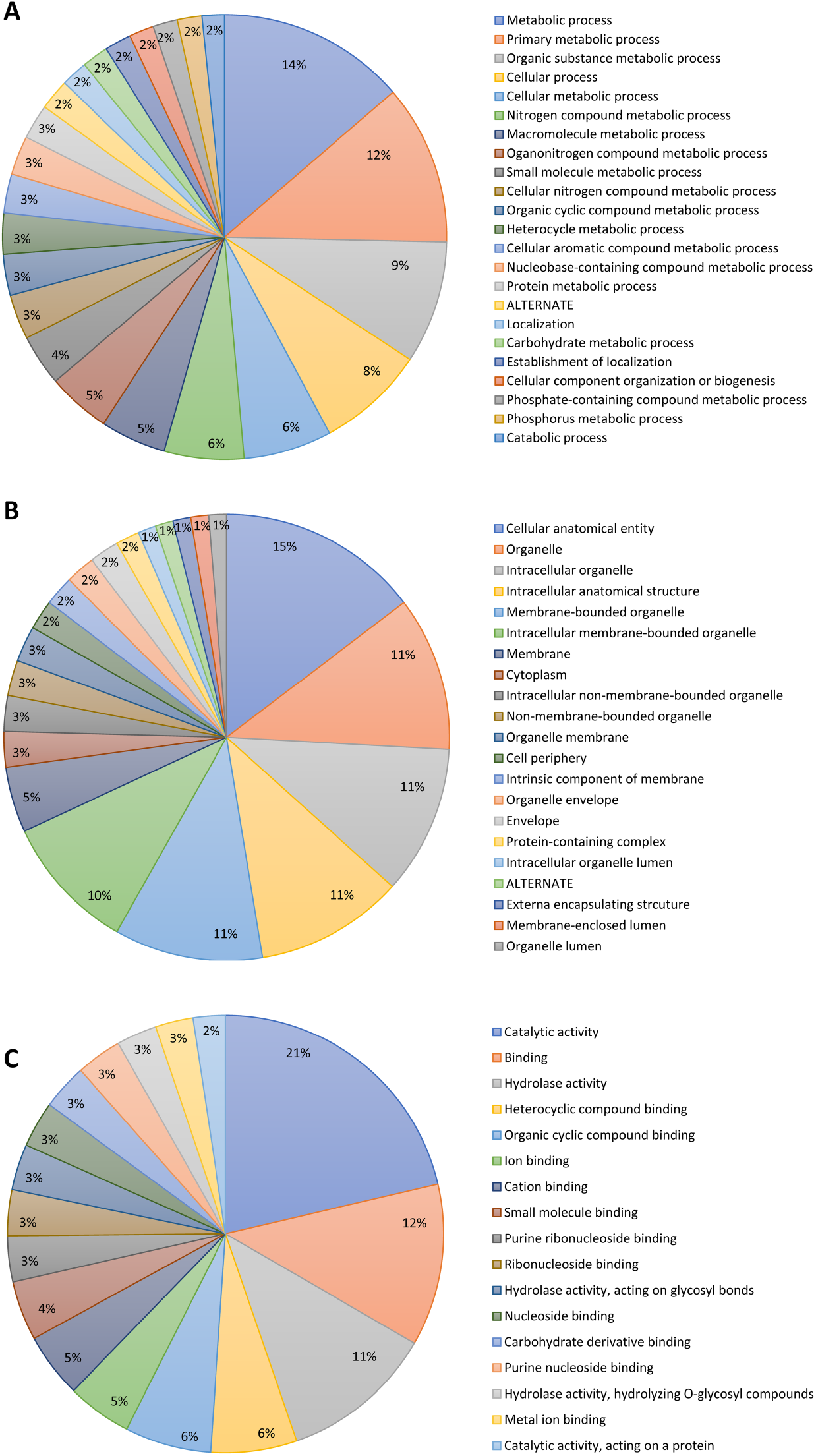
Functional annotation and analysis with OmicsBox of the proteins identified in *N. crassa* EVs. A) Biological processes; B) Cellular components; and C) Molecular function.

## 4. Discussion

This work shows that *N. crassa* releases EVs during vegetative growth in VMM under laboratory conditions. Previous reports in other fungal species have identified EVs heterogeneous in size, ranging from 10 to 1000 nm (Bielska and May, 2019). Size estimation of EVs has been obtained with techniques such as DLS, nanoparticle tracking analysis (NTA), or electron microscopy (Piffer et al., 2021). NTA allows to separate EVs size distributions at a higher resolution than DLS. NTA analyzes the trajectory of individual particles through their scattering, while DLS analyzes the intensity of the fluctuations of the light scattered by a set of particles. While DLS is not optimal to obtain accurate measurements of polydispersed suspensions, not all laboratories have access to an NTA technology.

There are previous publications where the method used for characterization of fungal EVs is DLS (Silva et al., 2014; Vargas et al., 2015), and it was further complemented by TEM analyses. We selected the number-based distribution from DLS results since it has been reported to have the highest concordance with TEM measurements (Bäuerl et al., 2020; Wooff et al., 2020). The EV measurements obtained with DLS in this work were within the range of previously reported values for other fungal EVs. In general, the average *d*_*h*_ values we obtained for *N. crassa* EVs by DLS were larger than the *d*_*p*_ values measured by TEM. On one hand, this difference in size could be due to intrinsic differences between the two methods, such as shown with polyethylene glycol or hydroxypropyl methylcellulose acetate succinate nanoparticles, which have a hydrated polymer corona shell that disappeared in a high-vacuum TEM environment (Wilson and Prud’homme, 2021). It is possible that the EVs of *N. crassa* present an external layer of proteins, which is not seen by TEM but is detected by DLS. A layer of mannoproteins was recently identified by Cryo-EM around EVs from *Cryptococcus neoformans, C. deneoformans*, and *C. deuterogattii* (Rizzo et al., 2021). On the other hand, the DLS approach assumes that particles have a sphere-like shape, even if they correspond to clusters of particles. Particle clusters can lead to enhanced scattering intensity (Dobbins and Megaridis, 1991), which would support the idea that the larger *d*_*h*_ values obtained by DLS correspond to the EV clusters that we observed by TEM. Both ultracentrifugation (Linares et al., 2015) and salt in the medium (Muschol and Rosenberger, 1995), which reduces EVs’ repulsive forces, could enhance EV aggregation.

For TEM imaging, areas with membranous-like particles of spherical shape were selected. We used two contrast agents: 1) osmium tetroxide vapors (Barland and Rojkind, 1966; Huang et al., 2020) as a selective agent for unsaturated lipids (Belazi et al., 2009), and 2) uranyl acetate, which is most commonly used to stain EV for TEM preparations in filamentous fungi (Bleackley et al., 2020; Hill and Solomon, 2020; Silva et al., 2014). Osmium tetroxide vapors gave similar results in terms of EV morphology and *d*_*p*_ in comparison to uranyl acetate. The use of vapors minimizes the presence of artifact-like vesicles that may be present in liquid solvents as impurities. In addition, it can be used simultaneously as fixative, reducing extra steps of primary fixation with aldehydes. Notably, osmium tetroxide vapors have been helpful in fixing delicate fungal structures such as conidia, conidiophores and aerial hyphae (Coetzee et al., 2005; Springer, 1989). Our results validated the use of osmium tetroxide vapors for EVs visualization.

In this study, we show evidence that *N. crassa* secretes nanosized EVs ranging from 24 to 140 nm, which fall within the reported sizes of exosomes. The average *d*_*p*_ values of EVs measured by TEM (34 or 38 nm depending on the staining method used) match the dimensions of ILVs contained inside MVBs in previous *N. crassa* reports (Rico-Ramírez et al., 2018; Roberson, 2020) and have similar sizes to the ones from *Ashbya gossypii* (Gibeaux et al., 2013), *Monoblepharis macrandra* (Dee et al., 2015), *Gilbertella persicaria* (Bourett et al., 2007), *Sclerotium rolfsii* (Roberson and Fuller, 1988), *Fusarium acuminatum* (Howard, 1981), and *Conidiobolus coronatus* (Fisher et al., 2018). This allows us to speculate that the EVs reported here could correspond to exosomes that have been released to the medium upon fusion of MVBs with the PM (Fig. 4). However, to date, there is no evidence of MVBs fusing to the PM in *N. crassa* or in any of the other fungal species mentioned above and further research is needed. Ongoing studies in *N. crassa* analyzing the subcellular localization of MVBs and the role of ESCRT components in MVBs biogenesis will help elucidate the route of synthesis of exosomal EVs. There are very few accounts by TEM of individual EVs localized in the periplasmatic space in *C. neoformans* (Wolf et al., 2014), *Magnaporthe grisea* (Bourett and Howard, 1996), *Botrytis cinerea* (Roberson et al., 2010), *Trichoderma reesei* (de Paula et al., 2019), and *N. crassa* (Roberson, 2020). Unfortunately, capturing rapid exocytosis events is challenging, even when using fast fixation procedures like cryo-fixation, limiting the amount of information available on exosomes and ectosomes *in situ*.

**Fig. 4.**
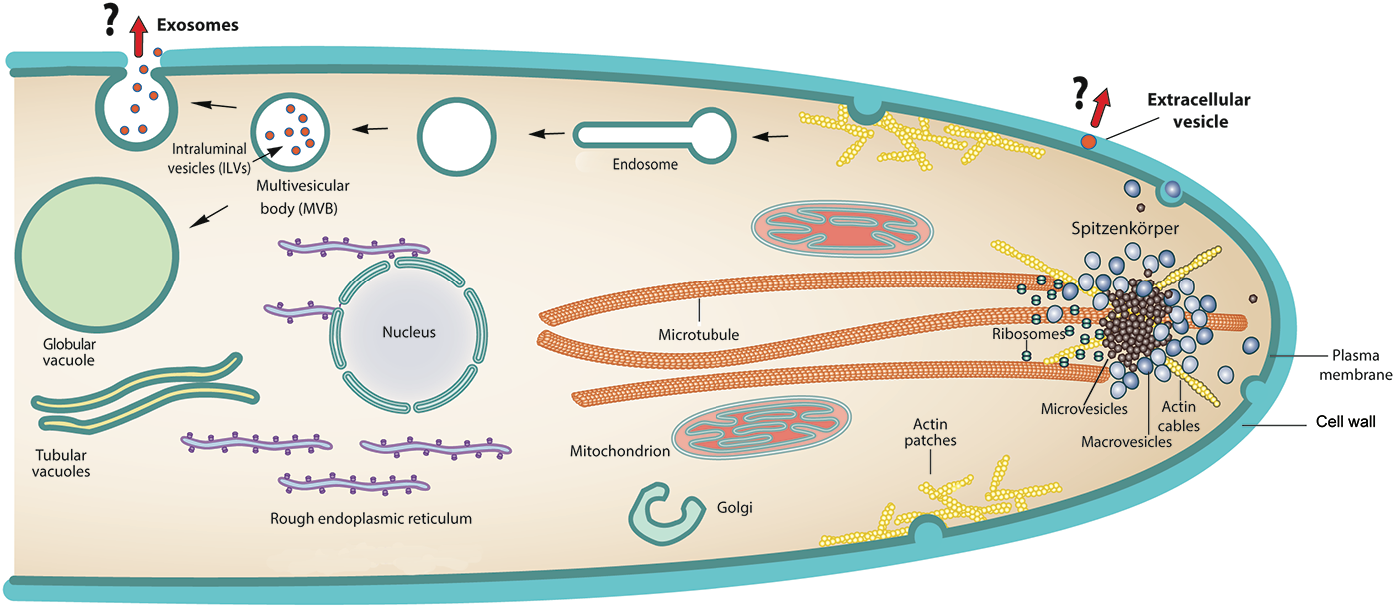
Schematic representation of the suggested secretory pathways leading to the biogenesis of vesicles in *N. crassa*. In subapical regions, exosomes would correspond to the ILVs (30-40 nm in diameter) released to the extracellular space upon fusion of MVBs with the PM. At the apex, secretory macrovesicles and microvesicles carrying cell wall synthesizing activities in a zymogenic state concentrate at the Spitzenkörper before fusing with the PM. Extracellular vesicles of unknown origin are found occasionally within the apical cell wall.

Proteomic analysis of fungal EVs has allowed identification of up to thousands of proteins, although most reports detect in the order of hundreds and some even fewer than 100 proteins. The results depend primarily on the mass spectrometry technology used (Bleackley et al., 2019; Rizzo et al., 2020; Sanderson et al., 2019). We identified 252 proteins in *N. crassa* EVs. The genome of *N. crassa* contains orthologues of exosomal marker proteins commonly reported in animal cells: two sequences encoding tetraspanins (PLS-1; NCU07432, TSP3; NCU09755), two annexins (ANXC4; NCU02139, ANX14; NCU04421), and one flotillin (FLOT; NCU05899) (Andreu and Yanez-Mo, 2014; Jeppesen et al., 2019; Khalaj et al., 2015). Nevertheless, none of these components were detected in these proteomic analyses, consistent with reports that indicate the lack of these proteins in fungal EVs (Zhao et al., 2019) and suggesting mechanisms for fungal EVs biogenesis different from those reported in mammalian cells.

The protein content of EVs can vary depending on the synthesis pathway, the cells’ physiological state, or the influence of the extracellular environment (Bielska and May, 2019). Enzymes involved in carbohydrate metabolism have been identified in EVs of pathogenic fungi, including *H. capsulatum* (Albuquerque et al., 2008), *Aspergillus fumigatus* (Souza et al., 2019), *C. albicans* (Vargas et al., 2015), and of the non-pathogenic fungus *T. reesei* (de Paula et al., 2019). In this work, we identified a glucoamylase, an amylase, a cellulase, and a high diversity of glycosidases. Glycosidases have been reported in exosomes surfaces and seem to be involved in the mobilization of growth factors and degradation of extracellular matrix macromolecules (Sanderson et al., 2019), which are mainly composed of polysaccharides. Diverse polysaccharides, including α-glucans (mainly α-1,3-glucan), β-glucans (β-1,6-branched β-1,3-glucan), galactomannan, and chitin are needed for a proper cell wall architecture in filamentous fungi (Yoshimi et al., 2017). Some glycosidases are responsible of the synthesis or degradation of these polysaccharides during cell wall remodeling. Moreover, the classical secretory process plays an essential role in cell wall expansion in filamentous fungi since secretory vesicles containing cell wall biosynthetic enzymes fuse with the PM (Peberdy, 1994). Surprisingly, cell wall biosynthetic enzymes that are predicted to be inserted into the PM, such as chitin synthase and glucan synthase, traditionally found in secretory vesicles, were also enriched in yeast EVs (Karkowska-Kuleta et al., 2020; Zhao et al., 2019). Enzymes related to chitin biosynthesis have also been found in EVs of *A. fumigatus* (Souza et al., 2019) and *T. reesei* (de Paula et al., 2019). In *N. crassa* no chitin or glucan synthases were identified in EVs, reinforcing previous studies, where chitin and *β*-1,3-glucan synthase were associated with the secretory microvesicles (chitosomes) and macrovesicles (Fig. 4), respectively (Sánchez-León et al., 2011; Verdin et al., 2009). Presumably, chitin or glucan synthase enzymes are incorporated into the PM since they are needed for *in situ* cell wall synthesis during polarized growth and are not secreted.

In addition, we identified other proteins associated to cell wall remodeling processes, including diverse 1,3-ß-glucanosyltransferases, anchored cell wall proteins (ACW), non-anchored cell wall proteins (NCW), GPI-anchored cell wall proteins (NCU09175, GE21DRAFT_1437 and GE21DRAFT_6010), CFEM-domain containing proteins (GE21DRAFT_5446, GE21DRAFT_1310533, and GE21DRAFT_8905), and a catalase. Glucanosyltransferases participate in cell wall remodeling and have been identified in hyphae and conidia of *N. crassa* (Ao et al., 2016). The glycoside hydrolase/glycosyltransferase BGT-2 (NCU09175) has been previously identified in the apical zone of *N. crassa* hyphae (Martínez-Núñez and Riquelme, 2015). Corresponding homologues have been previously reported in EVs of *C. albicans, S. cerevisiae, H. capsulatum*, and *T. reesei* (Adav et al., 2011; Albuquerque et al., 2008; Dawson et al., 2020). NCW and ACW proteins, other GPI-anchored cell wall proteins, and CFEM-domain containing proteins, play also essential roles in cell wall biosynthesis and remodeling during different stages of *N. crassa* development (Ao et al., 2016; Bowman and Free, 2006; Patel and Free, 2019). ACW-3, -4, -5, -8, -10, -11 and -12 detected here in *N. crassa* EVs, have been previously identified by proteomic analysis of the cell wall of vegetative hyphae and conidia of *N. crassa* (Ao et al., 2016; Maddi et al., 2009; Patel and Free, 2019). CFEM-domain containing proteins have also been localized in vegetative hyphae and conidia walls of *N. crassa* (Ao et al., 2016). CFEM (Common in several Fungal Extracellular Membrane) proteins have a GPI anchor domain and an eight-cysteine domain (Kulkarni et al., 2003). Some of the proteins that contain CFEM domains are known as anchored cell wall (ACW) proteins and play an essential role in cell wall remodeling at different stages of *N. crassa* development (Ao et al., 2016) or are involved in pathogenesis processes in *Magnaporthe grisea* (Kulkarni et al., 2005), *Botrytis cinerea* (Zhu et al., 2017) and *Candida albicans* (Pérez et al., 2011). A CFEM-domain-containing protein has been recently identified in *Fusarium oxysporum* EVs proteome, supporting a role for EVs in host-pathogen interactions (Garcia-Ceron et al., 2021). Finally, catalase 3 (GE21DRAFT_4671) has a role during oxidative stress and has been found in cell walls of vegetative hyphae and conidia of *N. crassa* (Maddi et al., 2009; Michán et al., 2002).

Of the total identified proteins of *N. crassa* EVs, the cell wall protein PhiA (NCU00399) and the clock-controlled gene protein CCG-14 (NCU07787) were the most abundant, representing the 45.46% of the emPAI total value. PhiA is an important protein for phialide formation during conidiation in *A. fumigatus* (Glaser et al., 2009) and *A. nidulans* (Melin et al., 2003). It also has a role in cell wall biosynthesis in *A. niger* when exposed to stress conditions (Schachtschabel et al., 2012). The second most abundant protein was CCG-14, whose expression is controlled by the circadian clock and is light sensitive (Cockrell et al., 2015). Clock-controlled genes have a critical role in the self-synchronization of cell cycles. They have been reported in exosomes from human and mouse cells (Tao and Guo, 2018). CCG-14 belongs to the superfamily of RlpA-like, with a cerato-platanin domain. It has a signal peptide region, predicted to be secreted. Members of the cerato-platanin family have a role in host-plant interaction in *Ceratocystis fimbriata* (Sbrana et al., 2007). CCG-14 is found in *N. crassa* cell wall of conidia and mycelia. This protein could act as a hydrophobin and an expansin due to its sequence similarity with cerato-platanin, a phytotoxin widespread in other ascomycetes (Krach et al., 2022). CCG-15 (NCU08936), known as AWC-1, was also identified in this study and has been found in the cell wall of both vegetative hyphae and conidia (Ao et al., 2016; Maddi et al., 2009). CCG-15 is highly similar to SPS2 of *S. cerevisiae* involved in sporulation (Zhu et al., 2017). CCG-15 belongs to the receptor L-domain superfamily. It has a signal peptide region and a C-terminal disordered region. In vesiclepedia, a community compendium for EVs, we found some genes related to circadian rhythms, such as a cryptochrome 1 (CRY1: http://microvesicles.org/gene_summary?gene_id=1407) and period genes PER1: http://microvesicles.org/gene_summary?gene_id=5187, PER2: http://microvesicles.org/gene_summary?gene_id=8864, PER3: http://microvesicles.org/gene_summary?gene_id=8863). Interestingly, circadian rhythm-related proteins FOTG_02405 and FOTG_15292 (two frequency/period clock proteins) were reported in EVs from *Fusarium oxysporum* sp. *vasinfectum* (Bleackley et al., 2020). We hypothesize that *N. crassa* could use EVs to regulate rhythm events, such as photoperiodic responses for cell synchronization. Remarkably, the well-described *N. crassa* circadian rhythm has an underlying molecular basis similar to those of higher eukaryotic organisms’ clocks (Dunlap, 1999; Dunlap and Loros, 2004; Heintzen and Liu, 2007). The existence of specific conserved clock-controlled proteins in mammalian and fungal EVs suggests an EV-mediated mechanism that allows them to adapt to daily changes.

Finally, we identified two proteins involved in late endosome vacuole transport and retrograde transport, endosome to Golgi: the vacuolar protein sorting-associated protein VPS-13 (GE21DRAFT_1332386) and vacuolar protein sorting-associated protein VPS-5 (NCU04137), respectively. The phosphatidylinositol 3-kinase VPS-34 (GE21DRAFT_2066) was identified as well. It catalyzes the formation of the signaling lipid phosphatidylinositol-3-phosphate and it is a key factor in the regulation of autophagy, endocytic trafficking, and vesicular transport in *S. cerevisiae* (Reidick et al., 2017). The vacuolar protein sorting-associated proteins (VPS) are part of Endosomal Sorting Complex Required for Transport (ESCRT) that performs the topologically unique membrane bending and scission reaction away from the cytoplasm (Schmidt and Teis, 2012). Vps5 forms a complex with Vps17 that regulates the recognition of endosomal membrane bending and contributes to the formation of tubules or vesicles on membranes (Nothwehr et al., 2000; Seaman et al., 1998). According to this, VPS-5 and another vacuolar protein sorting-associated protein (GE21DRAFT_1332386) could participate in the biogenesis or vesicular traffic. In addition to, we founded in the EV proteome a syntaxin protein (NCU07939). Syntaxins proteins are key in the current model of SNARE-mediated membrane fusion and in conjunction with other SNAREs proteins, syntaxins mediate vesicle fusion in diverse vesicular transport processes along the exocytic and the endocytic pathway (Teng et al., 2001). Besides a coatomer subunit beta (GE21DRAFT_6111) was identified, the coatomer is a complex that coats membrane-bound transport vesicles and it plays an essential role in retrograde Golgi-to-ER transport and retrieval of dilysine-tagged proteins back to the ER (Letourneur et al., 1994). This result suggest that intracellular transport of exocytosis vesicles take place into the fungus, and they are released into the extracellular space through the endoplasmic reticulum to the Golgi and then to the plasma membrane. The transport of EVs could be mediated by SNARE proteins, such as syntaxin (NCU07939), and a coatomer complex (GE21DRAFT_6111). The biogenesis of EVs of *N. crassa* suggests that vacuolar protein sorting-associated proteins as VPS-5 and GE21DRAFT_1332386 are involved. This is the first report which shows vacuolar protein sorting-associated proteins in EVs in fungi.

Altogether these results indicate a population of EVs heterogeneous in size containing secretory proteins that are involved mainly in cell wall remodeling, vesicular traffic, and in communication across hyphae during vegetative growth.

## CRediT author statement

**Elizabeth Medina-Castellanos:** Conceptualization, Methodology, Investigation, Formal analysis, Data Curation, Original draft preparation. **Daniel Salgado:** Methodology, Investigation, Formal analysis, Data Curation, Original draft preparation. **Juan Manuel Martínez-Andrade:** Methodology, Investigation, Formal analysis, Original draft preparation. **Ruben Darío Cadena-Nava:** Writing - review and editing, Funding acquisition. **Meritxell Riquelme:** Conceptualization, Formal analysis, Data curation, Writing-Reviewing and Editing, Supervision, Funding acquisition.

## Acknowledgments

We would like to thank Dr. Eduardo Callegari from the Proteomics Core Facility at the Sanford School of Medicine, of the University of South Dakota, USA, for technical support. We are grateful to the National Laboratory of Advanced Microscopy (LNMA) at the Centro de Investigación Científica y de Educación Superior de Ensenada (CICESE) and to the Centro de Nanociencias y Nanotecnología (CNyN) of the National Autonomous University of Mexico (UNAM) for letting us use their equipment.

## Data availability

The mass spectrometry proteomics dataset generated in this study has been deposited to the vesiclepedia database (http://www.microvesicles.org/index.html) with the vesiclepedia ID (information pending upon publication).

## Abbreviations

EVs: extracellullar vesicles
SPK: Spitzenkörper
TEM: transmission electron microscopy
PM: plasma membrane
DLS: dynamic light scattering
LC-MS/MS: liquid chromatography with tandem mass spectrometry

## Figure legends

**Fig. S1.**
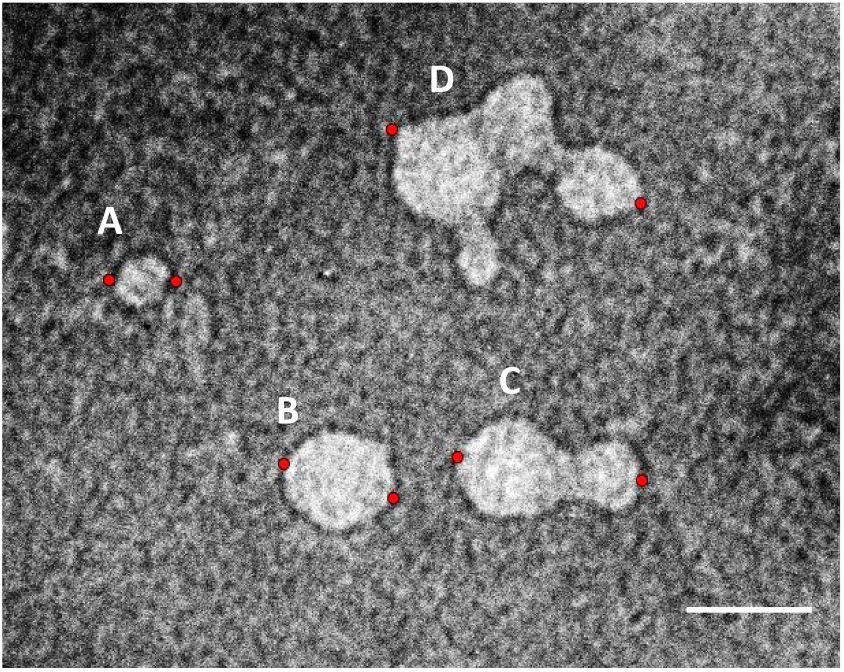
Comparison of clusters and individual EVs using osmium tetroxide vapors. A) Individual EV of 25 nm; B) Individual EV of 46 nm; C) Cluster of two EVs with a total diameter of 76 nm; and D) Cluster of more EVS with a diameter of 101 nm. The distance was determined between red points. Bar = 50 nm.

**Fig. S2.**
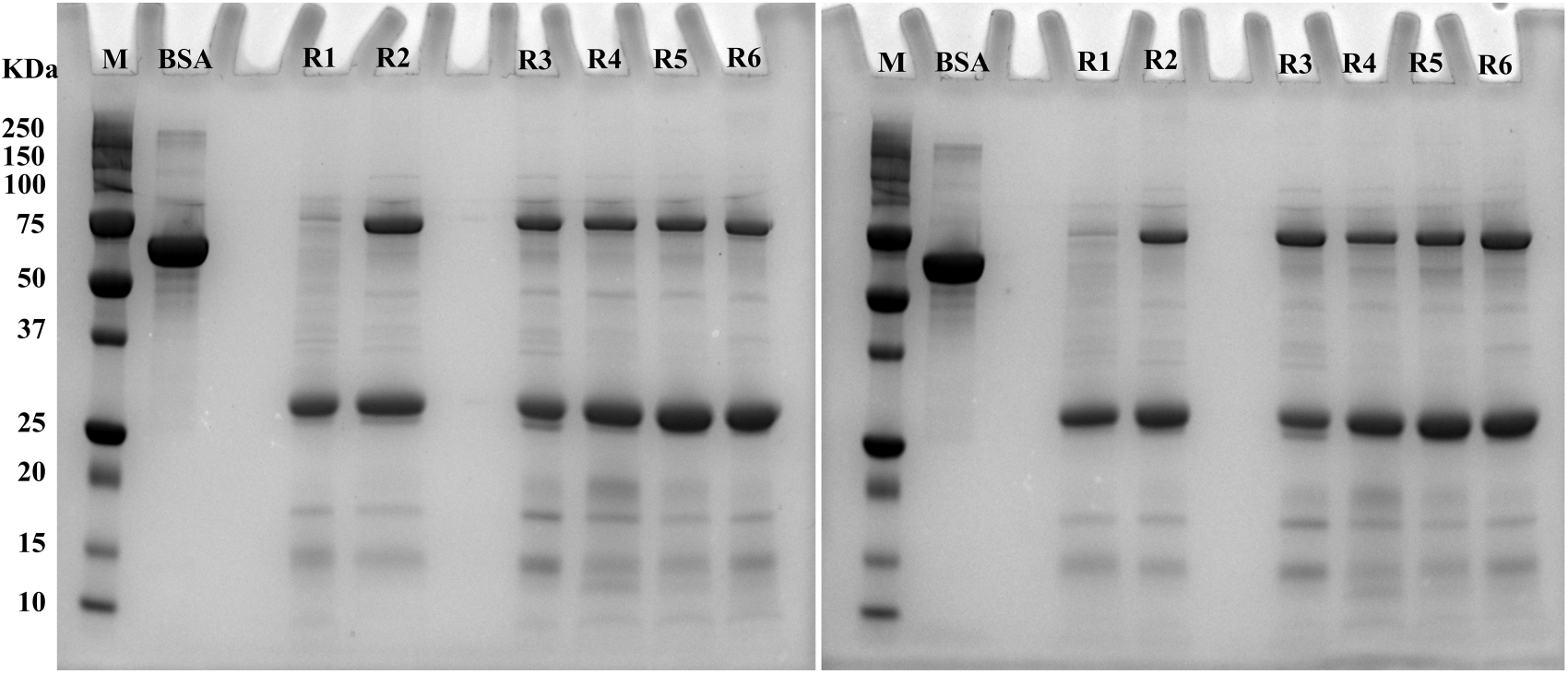
SDS-PAGE *N. crassa* EVs proteins. Each lane was loaded with 25 μg of protein. The gel was run at 250 V until the front line reached the end.

